# A geometric anthropomorphic phantom for quantitative susceptibility mapping: accuracy and repeatability

**DOI:** 10.64898/2026.09.01.748201

**Authors:** Padriac Hooper, Jin Jin, Kieran O’Brien, Markus Barth

## Abstract

Quantitative Susceptibility Mapping (QSM) relies on a tissue’s underlying macroscopic geometry to lead to measurable orientation-dependent field perturbations. To understand and assess QSM error in vivo, anthropomorphic phantoms provide a useful model that mimic the electromagnetic properties and morphology of underlying tissue.

Herein, we designed and manufactured an MRI compatible anthropomorphic phantom with cylindrical and spheroid compartments containing realistic susceptibilities to mimic hemorrhages, calcifications, and blood vessels. We estimated accuracy (ε, bias, RMSE) and repeatability (RC) of MEDI-susceptibility measurements within ROIs. We evaluated voxel-based agreement to validate susceptibility mapping under different acquisition conditions (3T versus 7T) and reconstruction algorithms (COSMOS versus MEDI).

Reliable MEDI-based susceptibility measurements were obtained from ellipsoids but not from straws. The ellipsoids (|ε| = 0.007 to 0.083 ppm at 3T; 0.050 to 0.118 ppm at 7T) were more accurate than the straws (|ε| = 0.084 to 0.190 ppm at 3T; 0.105 to 0.160 ppm at 7T). The repeatability coefficient across all 6 ROIs (RC = 0.652 ppm at 3T; 0.459 ppm at 7T) was substantially larger than across the 4 ellipsoid ROIs only (RC’ = 0.168 ppm at 3T; 0.141 ppm at 7T).

The accuracy at 3T (bias = -0.002 ppm, RMSE = 0.082 ppm) was better than the accuracy at 7T (bias = -0.056 ppm, RMSE = 0.092 ppm). Using voxels from the 4 ellipsoid ROIs, we observed excellent agreement between COSMOS and MEDI susceptibility maps at 3T, with linear regression of y=1.00x-0.01 (r=0.99). We observed some underestimation of MEDI susceptibility maps relative to COSMOS at 7T, with linear regression and y=0.93x-0.04 (r=0.99). The results imply that QSM reconstructions are reliable with 3T scanners but can be challenging with 7T scanners at high magnetic susceptibilities.

## 1. Introduction

Quantitative Susceptibility Mapping (QSM) is an advancing technique in MRI for quantifying the magnetic susceptibility of tissue (χ) from the phase component of T_2_*-weighted MRI data^1,2^. The magnetic susceptibility of tissue is sensitive to both tissue structure and composition and is used to identify various biomarkers within neuroimaging; iron^3,4^, blood deoxygenation^5,6^, and calcifications^7,8^. QSM is useful in the diagnosis and management of various neurological disorders, including neurodegenerative disease^3,9,10^, intracerebral hemorrhages^11^ and cancerous tissue^12^.

QSM involves various steps to process the phase (sometimes with the aid of magnitude) into a susceptibility map: phase unwrapping^13^, echo combination^14^, background field correction^15^ and dipole convolution^16–19^. The final step is particularly challenging due to the presence of zeros at spatial frequencies corresponding to a cone^20,21^; necessitating regularization within single-orientation QSM algorithms^22^, or by oversampling from multiple orientations to eliminate the null space within the kernel^23,24^. In addition, some susceptibility sources produce low SNR, making quantitative measurement challenging^25,26^. In QSM, it is impossible to define an absolute reference, and the choice of reference region has a statistical significance on clinical findings^27^.

Test objects (phantoms) provide a reference region of known susceptibility that make referencing viable, in contrast to *in vivo* QSM^28^. Phantoms can be designed to mimic the electromagnetic properties and morphology of tissue^29–31^ and provide a useful model to assess error within QSM reconstructions. They can provide a robust framework to assess systematic and random errors, free of patient-based experimental limitations^30^. Ideally, phantoms are composed of materials that match the desired imaging characteristics to function as a tissue surrogate^32^, and, human-like morphologies can be incorporated to replicate a realistic experiment^33^. It is appropriate to design a phantom containing compartments that resemble in vivo tissue at a macroscopic level. Within the brain, hemorrhages and microbleeds can roughly be regarded as spheroids, and blood vessels can be approximated as cylinders or ‘needle-like ellipsoids’^26,34^. In the literature, anthropomorphic phantoms have been used to assess MR signal properties^35,36^, thermal and electromagnetic properties^37–41^. However, these anthropomorphic phantoms contain structural components (e.g., nylon, ABS filament, etc.) that create artificial boundaries not usually observed *in vivo*; which leads to regions of low SNR, making quantitative imaging difficult^33,36,42^. 3D printing technology allows a broader range of phantom designs including more complex geometries, different materials, and compartments with well-defined boundaries. Since field perturbations are known to vary with geometry (e.g., cylinder versus spheroid)^26,34^, it is insightful to study the reliability of QSM reconstructions from within these region-of-interests. Moreover, the boundaries of geometric compartments are known beforehand, which means that potential errors can be easily identified and mitigated within QSM reconstructions.

Herein, we developed an anthropomorphic phantom with various geometric compartments containing realistic susceptibilities. Using well-defined region-of-interests (ROIs), we quantified the accuracy and repeatability of MEDI-based susceptibility measurements. We then evaluated voxel-based agreement to validate ill-conditioned single-orientation (MEDI) susceptibility maps relative to well-conditioned multi-orientation (COSMOS) susceptibility maps. We also evaluated voxel-based agreement between 3T and 7T scanners.

## 2. Methods

### 2.1. Design requirements

To create anatomically accurate MRI reference objects with features for quantitative analysis, the phantom requirements included the following: (a) encased in a human-sized skull-like object, scaled to the average head size of an adult male^43^; (b) intentional susceptibility features representing hemorrhages/microbleeds (spheroids) and blood vessels (cylinders); (c) To mimic realistic B_0_ fields, the nasal sinus cavities can be sealed off with air^37^. Extrusion-based printing was used to produce thick-walled structural pieces (skull, head), resin-based printing for thin-walled spheroid structures, and a straw with an adapter piece was used for thin-walled cylindrical structures.

### 2.2. CAD model

Annotated CAD designs of the realistically sized head phantom, spheroids, and the straw adapter were provided in Figure 1. All CAD designs and changes to other models were made using Inventor 2025 (Autodesk Inc., San Francisco, USA), requiring MeshEnabler and ThreadModeller plugins. Massachusetts General Hospital (MGH) has provided the CAD model of a head phantom with realistic B_0_ ^37^, which accomplishes the earlier stages of the anthropomorphic phantom model (human acquisition, segmentation, and surface creation), as described in MGH web page for ‘Angel 001’ head phantom (phantoms.martinos.org/MGH_Angel_001). The inner compartment is a skull, 3D printed as two matching parts, with many complex details such as the air cavities that can be sealed for realistic B_0_ field. The CAD design was edited to include sliding grooves for assembly between the inner and outer compartments to secure the phantom in place. The MGH ‘Angel 002’ head phantom (phantoms.martinos.org/MGH_Angel_002) incorporates a fill port into the brain volume, but has a prismatic volume, which was modified into a cylindrical volume that could be machined into an M10 screw fitting. A seal was produced using an appropriately sized O-ring (Freudenberg Sealing Technologies, Brendale, Australia). The outer posterior container was modified to have 2 plug-shaped fill ports for the removal of large air pockets that occur at narrow spaces. The phantom was scaled to a factor of 0.9:1 to be compatible within the radiofrequency coil (discussed later). The spheroids had three circular extrusions positioned 120^◦^ to one another about its central axis. Each had an M4 × 3 mm thread for bolt connection to the skull surface. At its enclosure was a fill port consisting of a thick neck (10 mm) and an M5 × 6 mm thread to provide a water-tight seal facilitated by a polytetrafluoroethylene (PTFE) (RS Group PLC, London, UK) tape layer. Each spheroid had an internal volume given by V = 4/3· π ·(15.3 mm)(12.3 mm)^2^ = 9.7 mL; the main cavity wall had a thickness of 0.7 mm. For the cylinders, a straw (6 mm diameter, 0.05 mm thickness, AGC-2, Quantum Design Inc, San Diego, USA) was placed at the end of a shaft (5.8 mm diameter). A drawing of a straw adapter was outlined, with the objective of holding each straw end rigid and consisting of two separate pieces: an insert and a collar. The insert and collar were designed to produce an interference fit, with an M2 × 3 mm thread for tightening the straw. Each insert had an M4 × 10 mm thread to provide a bolt connection to the skull surface with a water-tight seal. Each cylinder had an internal volume given by V = π ·(3mm)^2^ ·35mm = 0.99 mL; the wall thickness was 0.1 mm.

**Figure 1.**
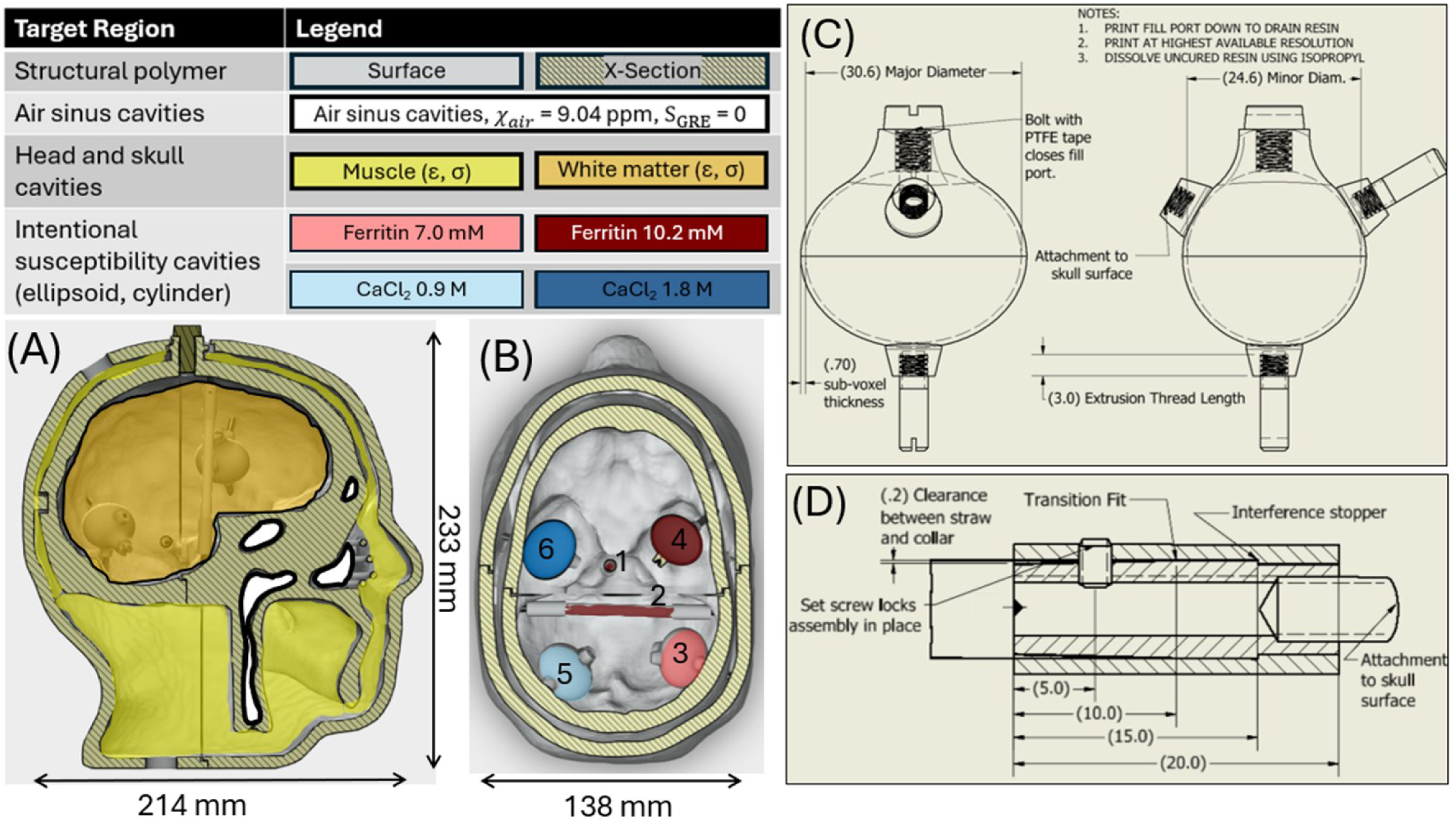
Head phantom CAD design. (A) Right side cross-sectional view indicating the bulk filling of the head, skull and air sinus cavities. (B) Top cross-sectional view revealing the positioning of intentional susceptibility features (spheroids, cylinders). (C) Front view of spheroid indicating its sub-voxel wall design, fill port, and M4 attachments. Major diameter (left, horizontal): 30.6 mm, minor diameter (right, horizontal & into the page): 24.6 mm, wall thickness: 0.7 mm, filling volume: 9.7 mL. (D) Detail view of straw adapter enclosure, which is mirrored at either end. The straw was slid over the adapter from a clearance fit (0.2 mm) to a transition fit (0 mm). An M2 bolt was threaded into the collar, the straw, then the shaft. An M4 bolt with PTFE tape was threaded at the opposing end to be attached to the skull surface. The filling volume of the straw was 0.99 mL (perpendicular orientation) and 1.56 mL (parallel orientation). The numberings in B indicated the ROI indexing used throughout the manuscript.

### 2.3. 3D-printing and treatment

A simplistic overview of anthropomorphic head phantom production is provided in Figure 2. The outer container was an assembly of 4 human-sized head parts (skull: anterior/posterior, head: anterior/posterior). 3D-printing of the outer container and straw adapter were accomplished with an UltiMaker S5 Pro Bundle (Ultimaker BV, Utrecht, Netherlands) using white acrylonitrile butadiene styrene (ABS) filament (0.1 mm layer height, 50% fill and triangle pattern) with dissolvable polyvinyl alcohol (PVA) supports. For the spheroids, the Figure 4 standalone stereolithography printer (3D System Inc, Rock Hill, USA) was used with medical amber resin at 30-micron (0.03 mm) layer height. The bulk of the uncured resin was drained from the hollow spheroid by 3D-printing the spheroid with the fill port facing down. The prints were then washed for five minutes each in two separate baths of isopropyl alcohol (to remove all uncured resin), then air dried for 60 minutes, followed by 90 minutes at 60 ^◦^C in a dual wavelength ultraviolet light box. After they came out of the ultraviolet light curer, the supports were sanded gently to remove 3D-printing supports. The spheroids were oven cured at 125 ^◦^C for approximately 30 minutes.

**Figure 2.**
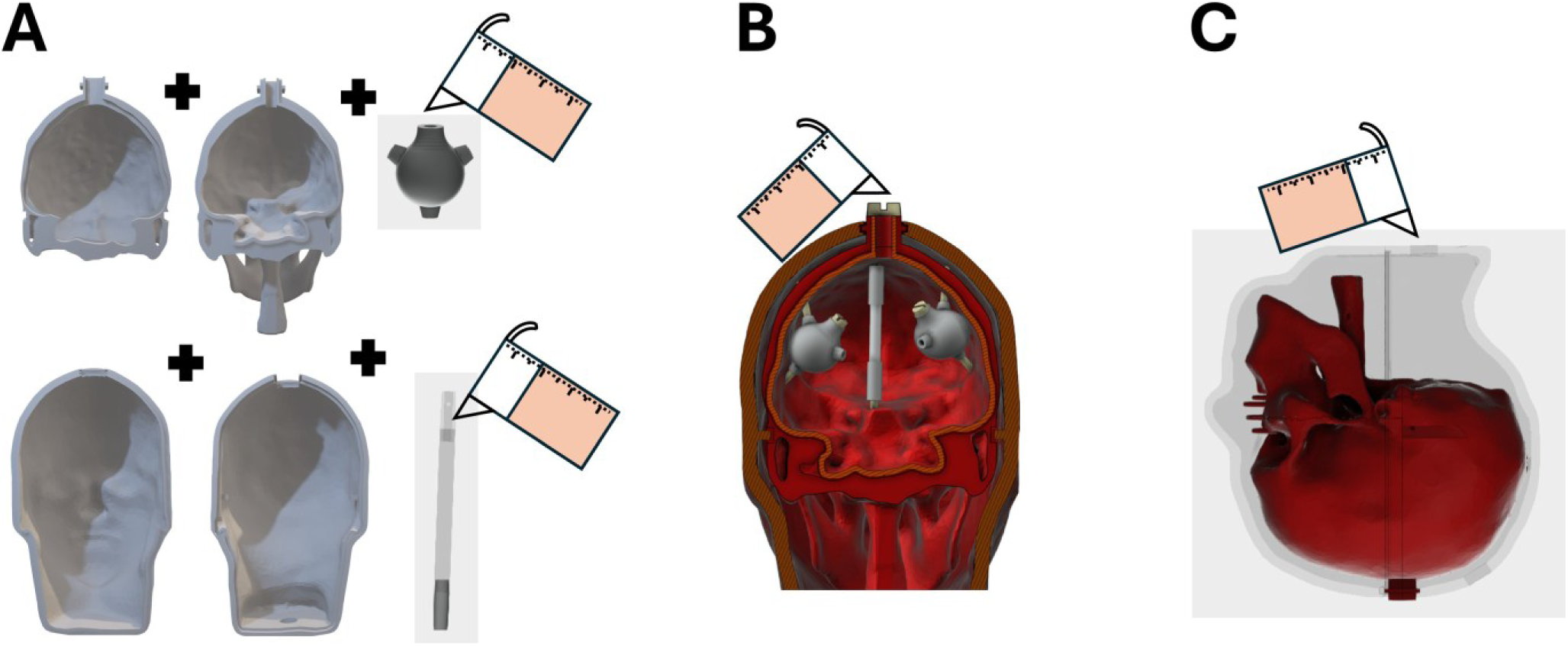
Simplistic overview of anthropomorphic head phantom production. A, 3D-printing and filling of geometric compartments and assembly into an anthropomorphic container, B, filling of brain cavity, C, filling of head cavity (shown in dark grey).

### 2.4. Phantom configuration, filling, sealing and assembly

Non-ferromagnetic doping materials: ferritin (paramagnetic) and CaCl_2_ (diamagnetic), which are theoretically susceptibility invariant with field strength^44^ were used. Iron-doped gel samples were prepared at 2 concentrations, ferritin: 7.0 and 10.2 mM Fe. Similarly, calcium-doped gel samples were prepared at 2 concentrations, CaCl_2_: 0.9 and 1.8 M. Each spheroid was filled with 1 of 4 gel preparations. The straws were filled with ferritin 10.2 mM Fe, oriented either parallel or perpendicular to B_0_ at the neutral frame. The ellipsoid fill port was sealed by white PTFE tape wrapped about a M5 nylon bolt, threaded into the fill port gently. The straws were connected to an adapter assembly, which was sealed by wrapping PTFE tape about each nylon bolt and set screw. The design and configuration of these geometric features is shown in Figure 1.

To fill the brain and head filling, polyvinylpyrrolidone-40 (PVP)-NaCl mixtures were prepared, following a protocol developed by researchers from New York University^45^. The skull was filled gradually (estimated volume: 1.05 L) and then allowed 48 h to settle. After waiting 48 h, the PVP gel mixture was carefully funneled over the brim of the fill port and then shaken to remove air bubbles at the fill port. The fill port was sealed by an O-ring (67008256, Freudenberg Sealing Technologies GmbH, Weinheim, Germany) about the shank and below the head of a M10 bolt (all bolts were nylon and purchased from RS Group PLC, London, UK). The head was filled gradually (estimated volume: 1.45 L) using both top and bottom plugs and then given 48 h to settle. The plugs to the outer cavities were sealed with Araldite 2020 epoxy adhesive (Huntsman Advanced Materials, Texas USA) Anthropomorphic phantom assembly (intermediate and final assembly) was shown in Figure 3. Araldite 2020 epoxy adhesive was applied sparingly for additional support at the nylon bolt attachment points. Air pockets were observed at the narrow space around the spheroid neck due to the contraction of the agarose gel after cooling.

**Figure 3.**
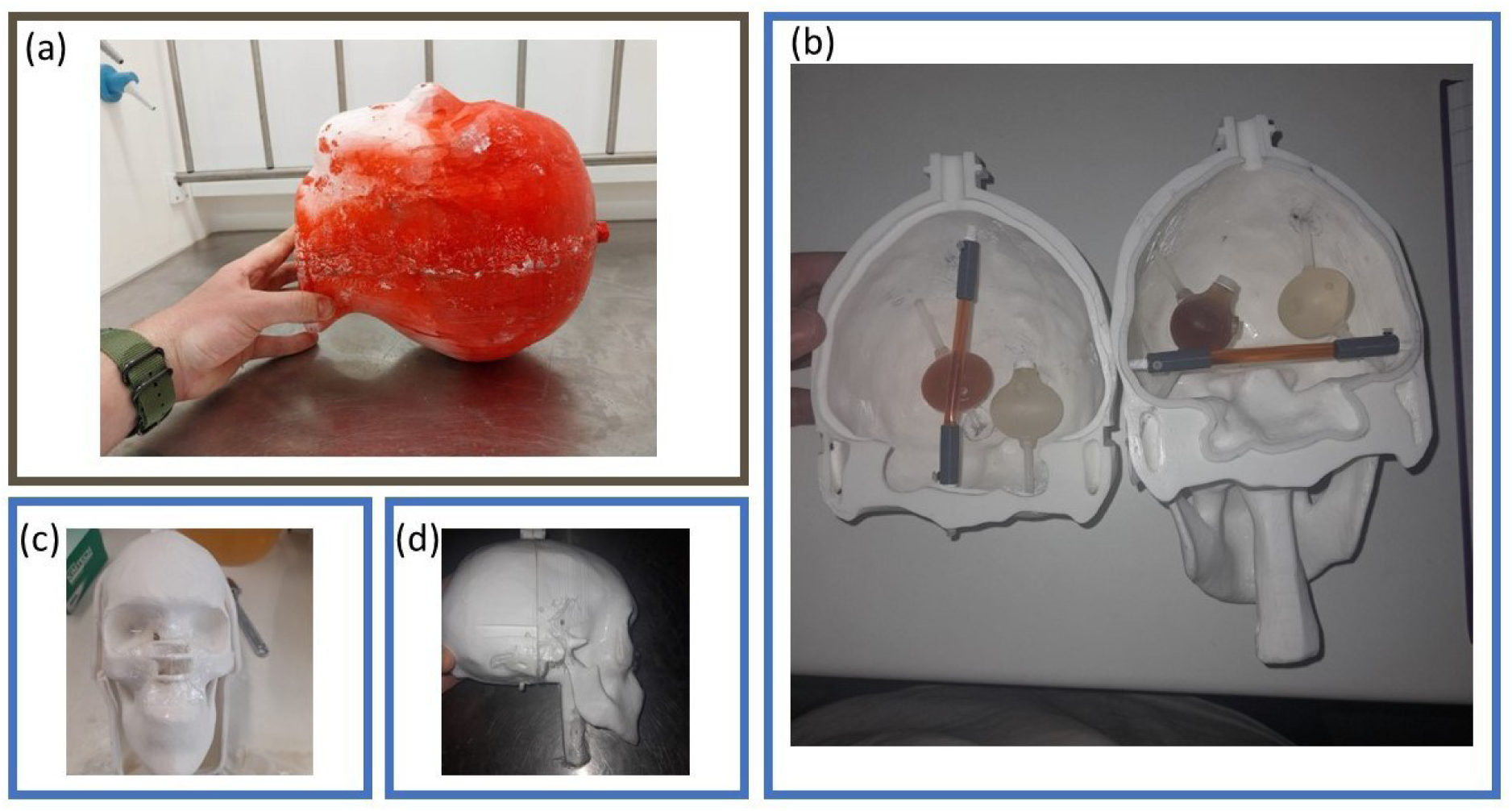
Anthropomorphic phantom assembly. A, The phantom fully assembled with red rubber coating applied for additional seal protection. B, Skull halves containing internal compartments. C-D, Seran wrap pressed and sealed with hot glue to isolate the skull air cavities from the gel.

**Figure 4.**
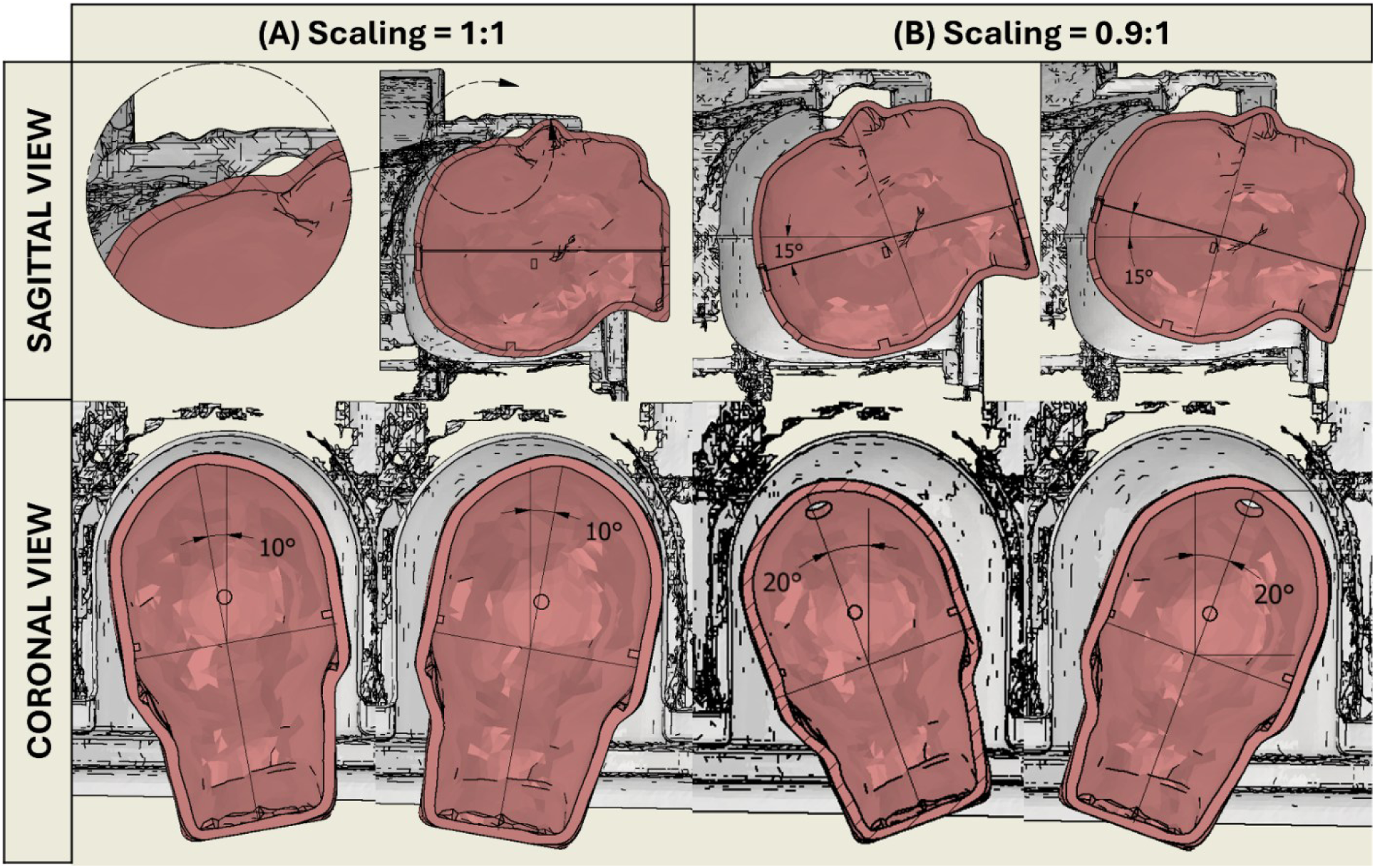
Rotations about the x- and y-axes of the scanner within the Nova Medical 32-Rx head coil used at 7 T, showing head sizes indicative of (A) a large male (scaling factor of 1:1) and (B) an average male (0.9:1 scaling factor). The sagittal view is perpendicular to the scanner’s x-axis (showing achievable x-axis rotations, θ), and the coronal view is perpendicular to the scanner’s y-axis (showing achievable y-axis rotations, ψ). The sagittal view detail showed that scaling factor of 1:1 cannot fit within the head coil.

### 2.5. Phantom scaling

To assess phantom scaling for 7 T MRI scanner (Siemens Healthineers, Forchheim, Germany), CAD assemblies were made using Inventor 2025 (Autodesk Inc., San Francisco, USA). Each CAD assembly consisted of the MGH head phantom CAD model at either 1:1 or 0.9:1 scaling; positioned within the 32 Rx Nova Medical head coil (the tightest fitting head coil intended for use). Then, using the CAD assembly as a reference, the maximum achievable rotations about the x- and y-axes of the scanner (θ, ψ) were estimated in 5° increments, then exported as a CAD drawing. Figure 4 showed the scaled phantom within the 32-Rx head coil, and the maximum achievable axial rotations. At 1:1 scaling, the phantom could not fit within the head coil (Figure 4, sagittal detail). A 0.9:1 scaling factor represented the largest phantom size that permitted sufficient rotational freedom while minimizing deviations from anatomical dimensions. At this scale, the phantom could rotate ± 15° about the x-axis and ± 20° about the y-axis, corresponding to 5 orientation sets: (θ, ψ) = (0°, 0°), (15°, 0°), (-15°, 0°), (0°, 20°) and (0°, -20°). The condition number (κ) provides a measure of conditioning for a non-square but full-ranked column vector, and is given below:

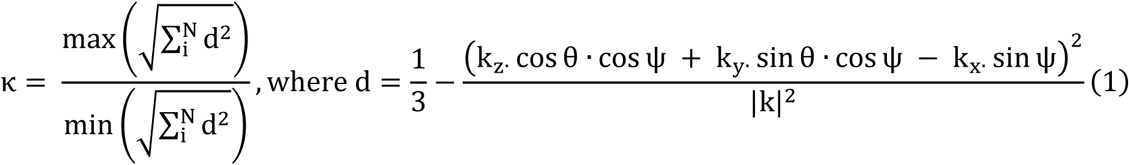

Where N is total number of orientations and d is the dipole kernel due to the projection of the k-space vector at each orientation frame *i* onto the B_0_ vector ([0, 0, 1]^T^). Combining the 5 orientations at [280, 320, 256] matrix size gives κ = 2.6, a well-conditioned reconstruction problem.

### 2.6. MR acquisition

The phantom was placed within the receive coil at the following configurations: a standard non-rotated acquisition, (θ, ψ) = (0°, 0°) and at least two rotations about the x- and y-axes, 6 acquisitions in total, as indicated in Supplementary Tables 1 & 2. We were unable to orient the phantom within the coil arrangement to a high degree of accuracy, as it has been done in a previous study^46^. Non-local field effects, especially in the B_0_ direction, leads to strong dependence of QSM on spatial coverage^47^. We mitigated these errors by using large FOVs (210 × 240 × 192 mm^3^) for MR acquisition. For each orientation, phase and magnitude images were acquired using a multi-echo gradient-recalled-echo (GRE) sequence, using parameters outlined in Table 1.

**Table 1.**
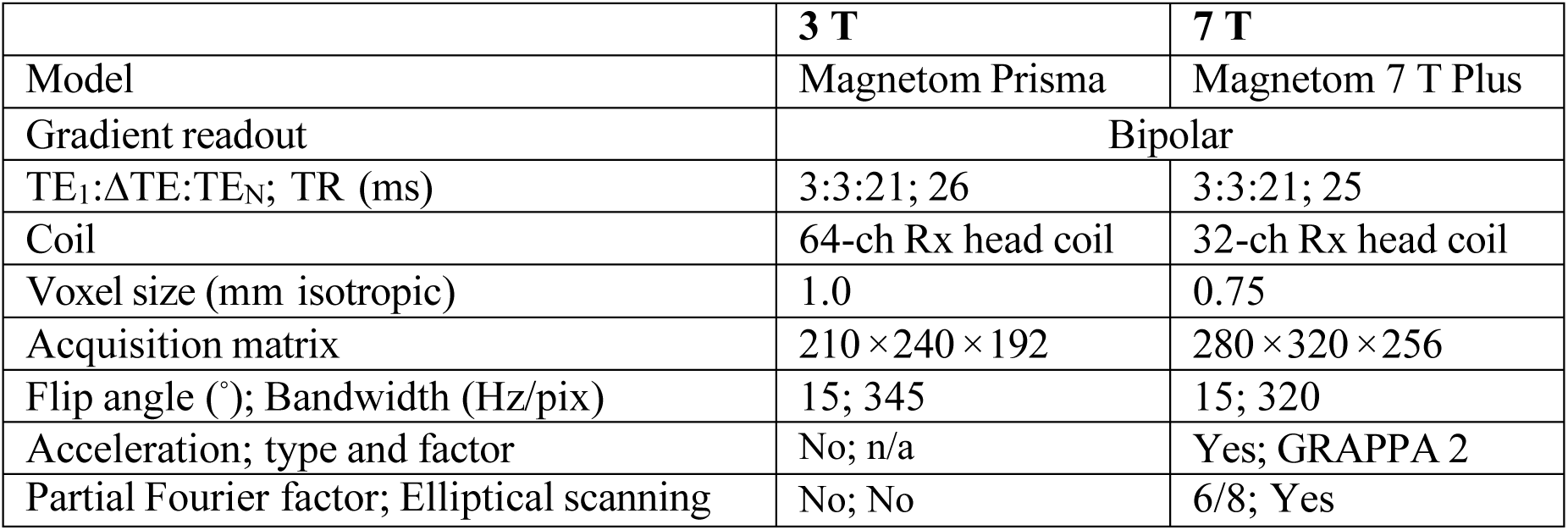
MR acquisition parameters used in the current study.

|  | 3 T | 7 T |
| --- | --- | --- |
| Model | Magnetom Prisma | Magnetom 7 T Plus |
| Gradient readout | Bipolar |  |
| TE<sub>1</sub>:ΔTE:TE<sub>N</sub>; TR (ms) | 3:3:21; 26 | 3:3:21; 25 |
| Coil | 64-ch Rx head coil | 32-ch Rx head coil |
| Voxel size (mm isotropic) | 1.0 | 0.75 |
| Acquisition matrix | $210 \times 240 \times 192$ | $280 \times 320 \times 256$ |
| Flip angle ($^\circ$); Bandwidth (Hz/pix) | 15; 345 | 15; 320 |
| Acceleration; type and factor | No; n/a | Yes; GRAPPA 2 |
| Partial Fourier factor; Elliptical scanning | No; No | 6/8; Yes |

### 2.7. Image pre-processing and corrections

A brain mask (M) was produced using FMRIB’s Brain Extraction Tool (BET; FSL 6.0.7)^48,49^ on the 1^st^-echo magnitude image, and holes introduced during BET were filled in. To reflect the faster dephasing and generally more rapid T_2_* decay at ultra-high-field, the final echo time used at 7T (9 ms) was set to 3/7 that used at 3T (21 ms). To improve the quality of the co-registrations in sections 2.9, 2.11, N4 bias field correction^50^ was applied to the magnitude images, then recombined into complex GRE data. To improve the SNR, MP-PCA denoising^51^ was applied to the complex GRE data. The image matrices were zero-padded (321 × 321 × 321 voxel^3^) to reduce aliasing associated with the Fourier transform. Phase offsets were corrected using MCPC-3D-S^52^. Geometric distortions were removed by generating a voxel displacement map from the odd and even field maps^53,54^, then unwarping the complex GRE data^55^ with tri-linear interpolation. A noise map was generated using the MEDI complex nonlinear fitting algorithm^56^. To compute the initial weights (W_0_): the noise map was inverted and normalized using the median and upper interquartile range (IQR), re-centered to 1, then outliers (defined as median + 3·IQR) were replaced with a 3 × 3 × 3 voxel box filtered copy. The relative residual (r) of measured complex GRE data S_measured_ (Equation 2) and simulated complex GRE data S_simulated_ (Equation 3) was computed as follows^57^,

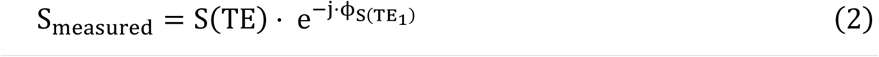

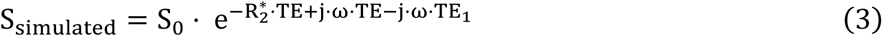

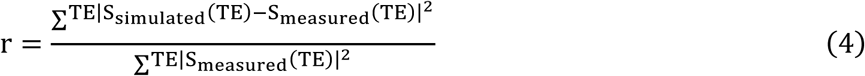

In these equations, the angular frequency (ω) was computed during field mapping, the relaxation rate (R_2_*) from the multi-echo magnitude data using ARLO^58^ and the proton density (S_0_) by extrapolating signal intensity to TE=0. The relative residual map was brought into a weighting component using a threshold, r_th_ = median(r) + 3·IQR(r), to derive the weighting map (W)^57^,

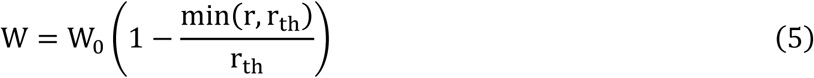

The multi-echo phase was combined using the MEDI complex nonlinear fitting algorithm^56^, then unwrapped using SEGUE^59^. We then eroded the brain mask using a 3 mm radius spherical structuring element. Background fields were corrected using PDF^60^, using the default parameters within the MEDI toolbox. The image matrices were restored back to the original FOV (210 × 240 × 192 mm^3^).

### 2.8. Single-orientation QSM

MEDI^16–18,56^ was performed for single-orientation QSM,

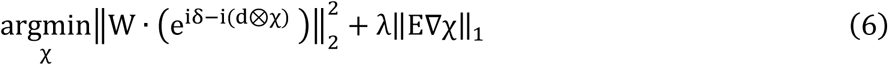

In which δ and d denoted local fields and dipole kernel, respectively. In the term λ‖E∇χ‖_1_, ∇ denoted the gradient operator, and E the edge weighting mask derived from the gradient magnitude image (∇m) by considering a given ratio of voxels (c∇=[0,3]) to be edges^16^:

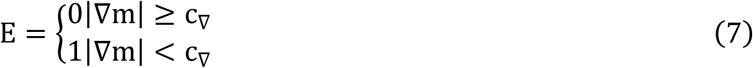

It was determined that c∇=0.3 provided a balance between morphological consistency and image fidelity (see Figure S1). MEDI was reconstructed with default regularization value: λ=1000.

### 2.9. Multi-orientation QSM

Transformation matrices were obtained by co-registering magnitude images from the *i^th^* frame to the neutral frame using FMRIB’s Linear Image Registration Tool^61^ (FLIRT; FSL 6.0.7) with six-parameter rigid transformations. The local fields (δ_i_) and weighting maps (W_i_) were then co-registered from the *i^th^* frame to a common reference frame with spline interpolation and normalized mutual information metric. COSMOS can be computed efficiently using a closed-form solution,

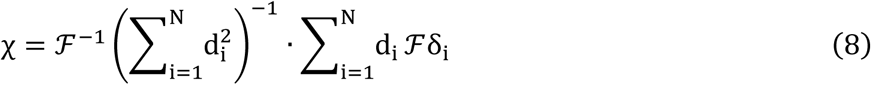

However, oversampling from multiple orientations cannot reconstruct susceptibility in signal void regions, such as those occupied by air. To eliminate solutions with sharp discontinuities at regions of high susceptibility and low SNR, we model a homogeneous susceptibility distribution by adopting image gradient regularization^62^,

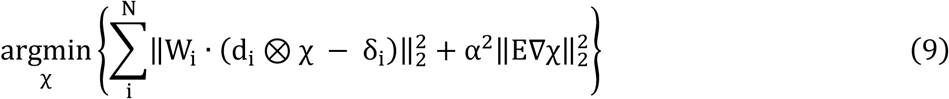

Equation (9) was solved using a conjugate gradient solver using the solution in Equation (8) as an initial guess, and edge weighting mask (E) was derived from the magnitude image acquired at the first orientation. The regularization parameter (α=1000) was stable over several orders of magnitude. A comparison of COSMOS-based susceptibility maps is provided in Table S3.

### 2.10. Referencing

Susceptibility maps were referenced to a 15 mm sphere positioned within the background filling (see Figure S2), and all reported susceptibility values are expressed relative to this reference.

### 2.11. Segmentation

Region-of-interest (ROI) segmentation was performed by performing the following operations:

- Removal of background filling (since doped agarose mixtures have a short T_2_* compared to filling fluid): R_2_* > 4 s^-1^.
- Removal of edge voxels: the L_2_-norm of the gradient field E was normalized by its global maximum, and voxels > 0.999 were classified as non-edges.
- Removal of unreliable voxels (from ellipsoids): r < median(r) + 1.5·IQR(r).
- Erosion of brain mask with a spherical structural element of 12 mm radius.

From the output mask, we analyzed connectivity (6-neighbor) and selected regions of size 200-10,000 mm^3^.

### 2.12. Statistical analysis

At each field strength, the MEDI susceptibility maps were registered to the common reference frame using the registration matrices derived in Section 2.9. For analyses in sections 2.11.2-2.11.3, the 2 straw ROIs (ROIs 1 and 2) were excluded.

#### 2.12.1. ROI accuracy, precision and repeatability

We reported the following ROI-based metrics:

- **Accuracy:** for each ROI, we defined “accuracy error” (ε) as the difference between QSM susceptibility and literature susceptibility (ε = χ - χ_lit_). χ_lit_ was computed by multiplying the literature molar susceptibility (76.6 ppm·M^-1^ for ferritin^63^ and -0.271 ppm·M^-1^ for CaCl_2_^44^) by molar concentration (indicated in Figure 1A). Across 6 ROIs, we computed bias= Σε and RMSE = √(Σε^2^_i_⋅n_i_ / Σn_i_), where n_i_ is the number of voxels within ROI *i*.
- **Precision & Repeatability:** for each ROI, we measured the within-region standard deviation (σ_R_) to assess precision. For each ROI, we measured the standard deviation across 6 acquisitions (σ_X_) to assess repeatability. We measured the repeatability coefficient (RC=2.77⋅√σ_X_^2^) across a range of ROIs (RC=all 6 ROIs, RC’=4 ellipsoid ROIs only).

#### 2.12.2. COSMOS-MEDI agreement

At each field strength, a “representative” MEDI susceptibility map was computed by taking the average over 6 acquisitions. For 3T and 7T susceptibility maps, a voxel-based linear regression and a Bland-Altman analysis was then performed to assess agreement between COSMOS and “representative” MEDI.

#### 2.12.3. Crossfield-strength agreement

To register the 7T data to the 3T data, we first obtained transformation matrices by co-registering the 7T reference magnitude image (resampled to 1 mm^3^) to the 3T reference magnitude image (already at 1 mm^3^) using FLIRT^61^ with six-parameter rigid transformations and normalized mutual information metric. We then co-registered the 7T COSMOS susceptibility map and the 7T “representative” MEDI susceptibility map (each resampled to 1 mm^3^) to the 3T common reference frame with spline interpolation. For COSMOS and “representative” MEDI susceptibility maps, a voxel-based linear regression and a Bland-Altman analysis were performed to assess agreement between 3T and 7T.

## 3. Results

### 3.1. Visual assessment

Figure 5 shows a visual assessment at both field strengths (3T, 7T), and reconstruction algorithms (MEDI, COSMOS), indicating how the construction materials appear in susceptibility maps, and how susceptibility contrast appeared within different shapes (ellipsoid versus straw). On ROI #2, the yellow arrowhead shows a strong paramagnetic susceptibility effect caused by the ABS (Δχ = 2.748 ppm)^64^. On ROI #5, the yellow arrowhead shows nylon (Δχ = -0.446 ppm)^42^ at the ellipsoid fill port. The green arrowhead shows a homogeneous susceptibility distribution within the ellipsoid marked as ROI #4, excluding its borders. The red arrowhead shows a heterogeneous susceptibility distribution within the straw marked as ROI #2.

**Figure 5.**
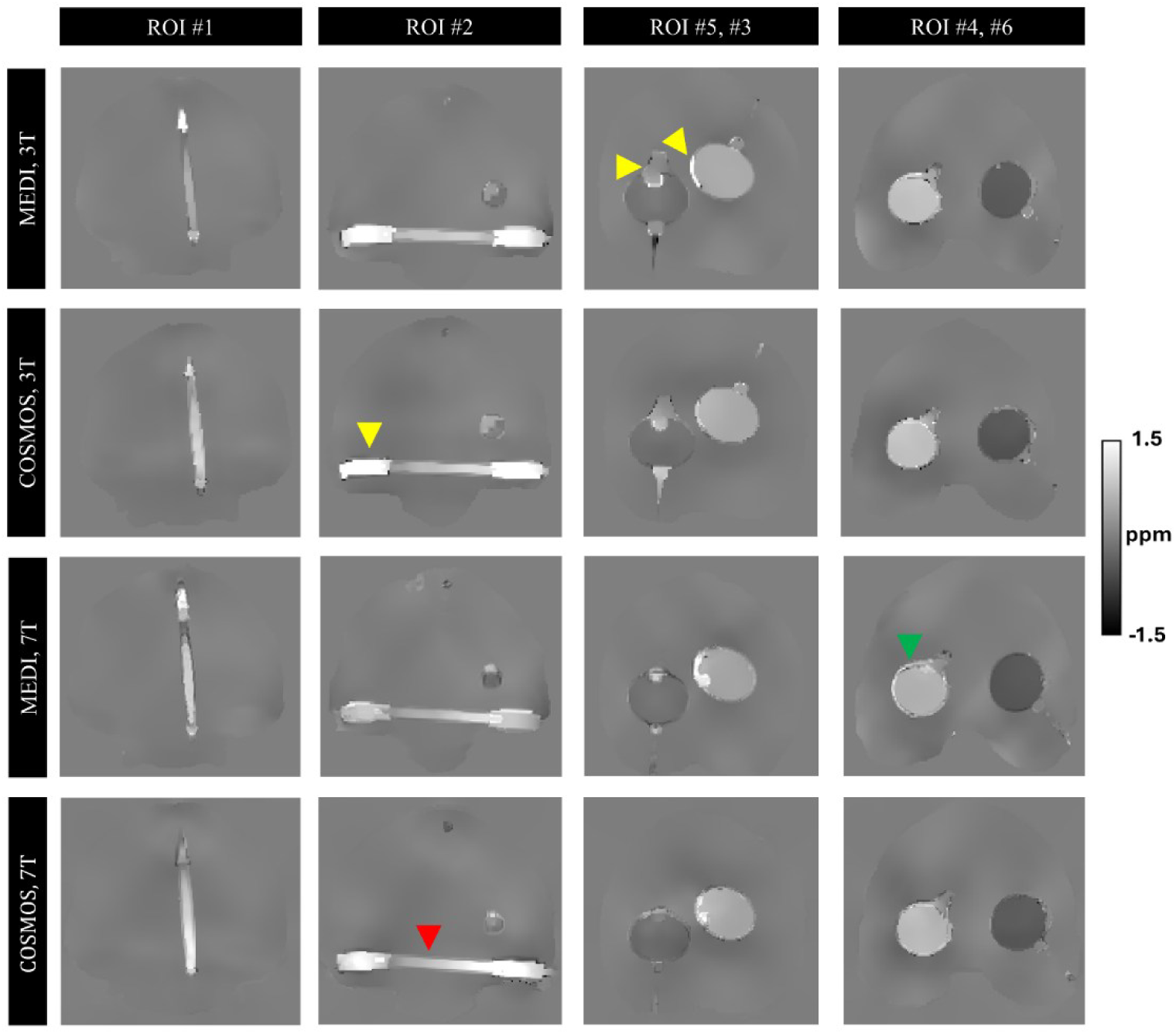
Visual assessment of susceptibility maps reconstructed with various algorithms (MEDI, COSMOS) and various field strengths (3T, 7T). On ROI #2, the yellow arrowhead shows a strong paramagnetic susceptibility effect caused by the ABS (Δχ = 2.748 ppm)^68^. On ROI #5, the yellow arrowhead shows nylon (Δχ = -0.446 ppm)^42^ at the ellipsoid fill port. The green arrowhead shows a homogeneous susceptibility distribution within the ellipsoid marked as ROI #4, excluding its borders. The red arrowhead shows a heterogeneous susceptibility distribution within the straw marked as ROI #2.

### 3.2. ROI-based quantitative comparison

The accuracy, precision and repeatability were summarized in Table 2. Please refer to Figure S3 for a scatter plot of QSM susceptibility measurements against literature susceptibility values (ROI accuracy), and within-region standard deviation (ROI precision) shown as error bars. For the paramagnetic ROIs (ROIs 1 to 4), ε<0 indicated susceptibility underestimation. Conversely, for the diamagnetic ROIs (ROIs 5 to 6), ε>0 indicated susceptibility underestimation. Except for ROI #1 at 3T, every QSM measurement was underestimated. The ellipsoids (|ε| = 0.007 to 0.083 ppm at 3T; 0.050 to 0.118 ppm at 7T) were more accurate than the straws (|ε| = 0.084 to 0.190 ppm at 3T; 0.105 to 0.160 ppm at 7T). The bias at 3T (-0.002 ppm) was lower than at 7T (-0.056 ppm). The RMSE at 3T (0.082 ppm) was lower than at 7T (0.092 ppm). The ellipsoids (σ_R_ = 0.013 to 0.036 ppm at 3T; 0.023 to 0.053 ppm at 7T) were more precise than the straws (σ_R_ = 0.206 to 0.295 ppm at 3T; 0.312 to 0.317 ppm at 7T). The repeatability of the ellipsoids (σ_X_ = 0.012 to 0.048 ppm at 3T; 0.012 to 0.039 ppm at 7T) was better than the repeatability of the straws (σ_X_ = 0.112 to 0.195 ppm at 3T; 0.083 to 0.134 ppm at 7T). The repeatability coefficient across all 6 ROIs (RC = 0.652 ppm at 3T; 0.459 ppm at 7T) was much greater than across the 4 ellipsoids (RC’ = 0.168 ppm at 3T; 0.141 ppm at 7T).

**Table 2.**
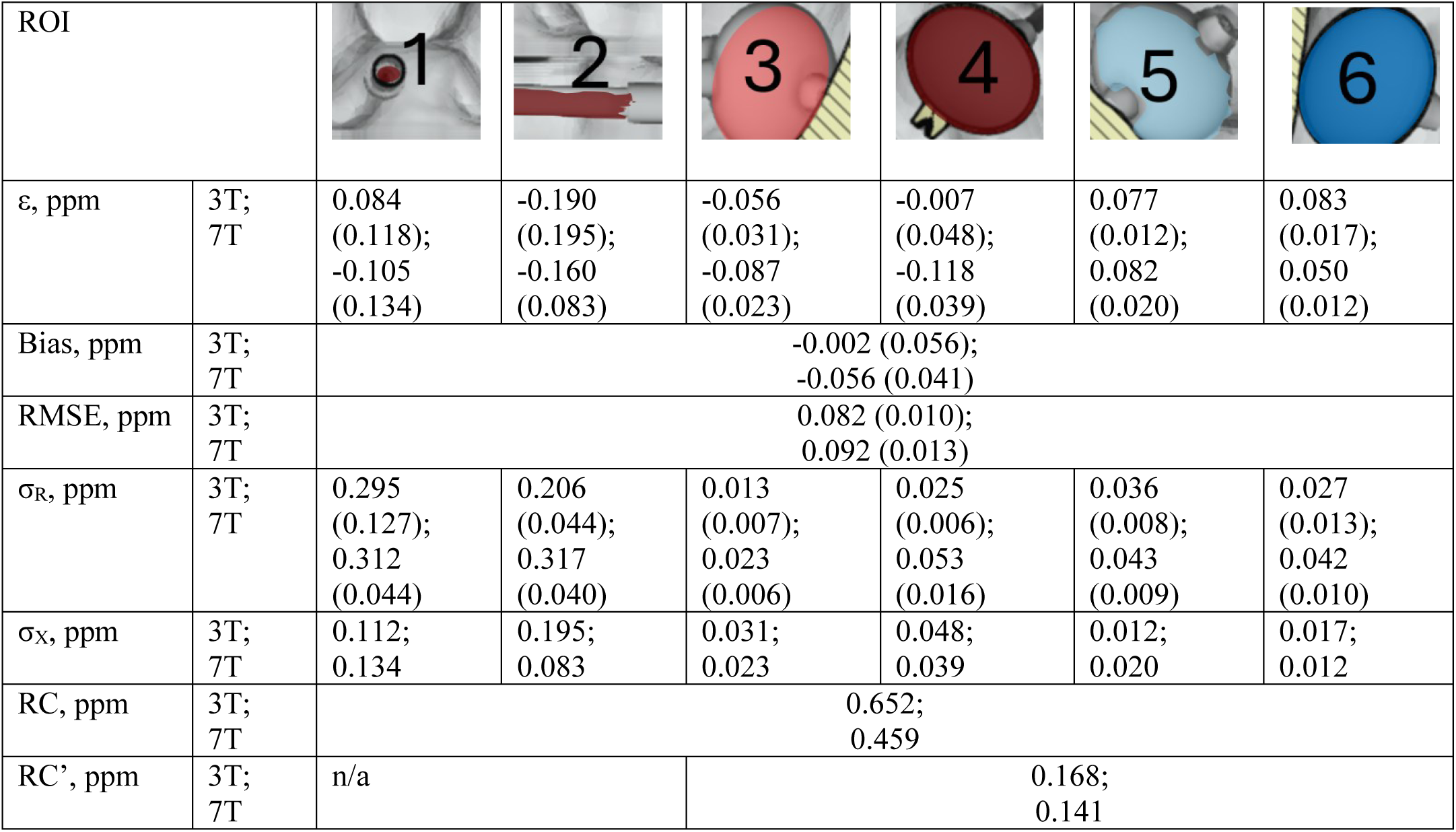
Accuracy, precision, and repeatability of ROI measurements. Accuracy error (ε=χ- χlit), bias, RMSE, and within-region standard deviation (σR) were reported as mean (SD) across 6 orientations. Repeatability metrics included cross-orientation standard deviation (σX), repeatability coefficients across all 6 ROIs (RC) and across 4 ellipsoid ROIs only (RC’).

### 3.3. COSMOS-MEDI agreement

For the 4 ellipsoid ROIs, Figure 6 provides a quantitative comparison between COSMOS and MEDI susceptibility maps. At 3T, COSMOS and MEDI were in close agreement. The mean difference was -0.008 ppm, and the 95% limit-of-agreement between -0.103 to 0.088 ppm. At 7T, MEDI underestimated COSMOS by 7%. The mean difference of -0.045 ppm, and the 95% limit-of-agreement between -0.187 to 0.098 ppm. In each case the correlation coefficient was 0.99.

**Figure 6.**
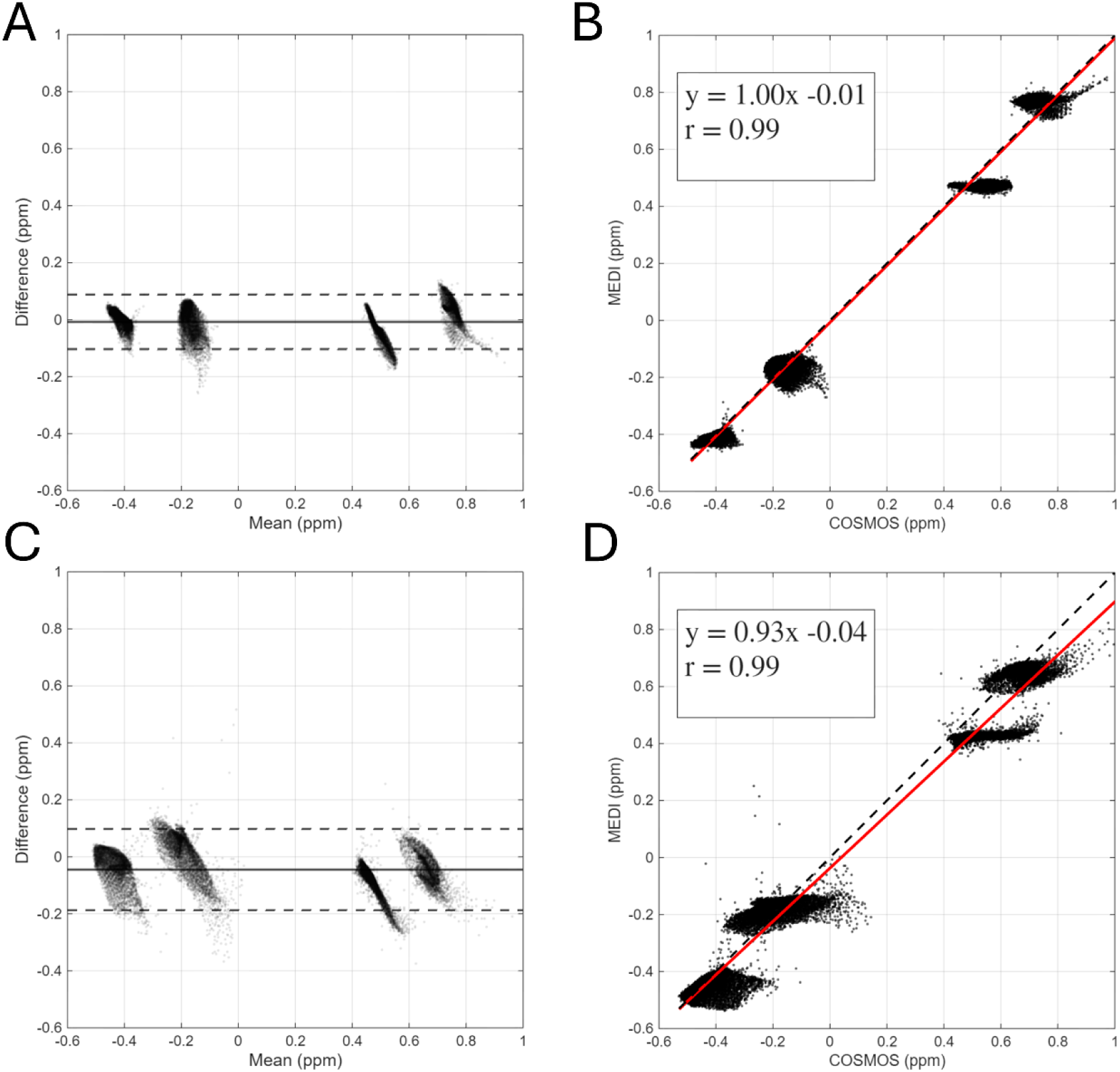
Quantitative comparison between COSMOS and MEDI at 3T (A-B) and 7T (C-D). For the Bland-Altman plots (A,C), the solid and dashed lines are the mean difference ± 2 × the standard deviation of the difference, respectively. For the correlation plots (B,D), the solid red lines indicate the trend lines in linear regression and the dashed line is the line of identity.

### 3.4. Crossfield strength agreement

For the 4 ellipsoid ROIs, Figure 7 provides a quantitative comparison between 3T and 7T. For MEDI susceptibility maps, 7T underestimated 3T by 4%. The mean difference was -0.030 ppm, and the 95% limit-of-agreement was -0.105 to 0.046 ppm. The correlation coefficient was 1.00. For COSMOS susceptibility maps, 3T and 7T were in close agreement. The mean difference was -0.014 ppm, and the 95% limit-of-agreement was -0.120 to 0.093 ppm. The correlation coefficient was 0.99.

**Figure 7.**
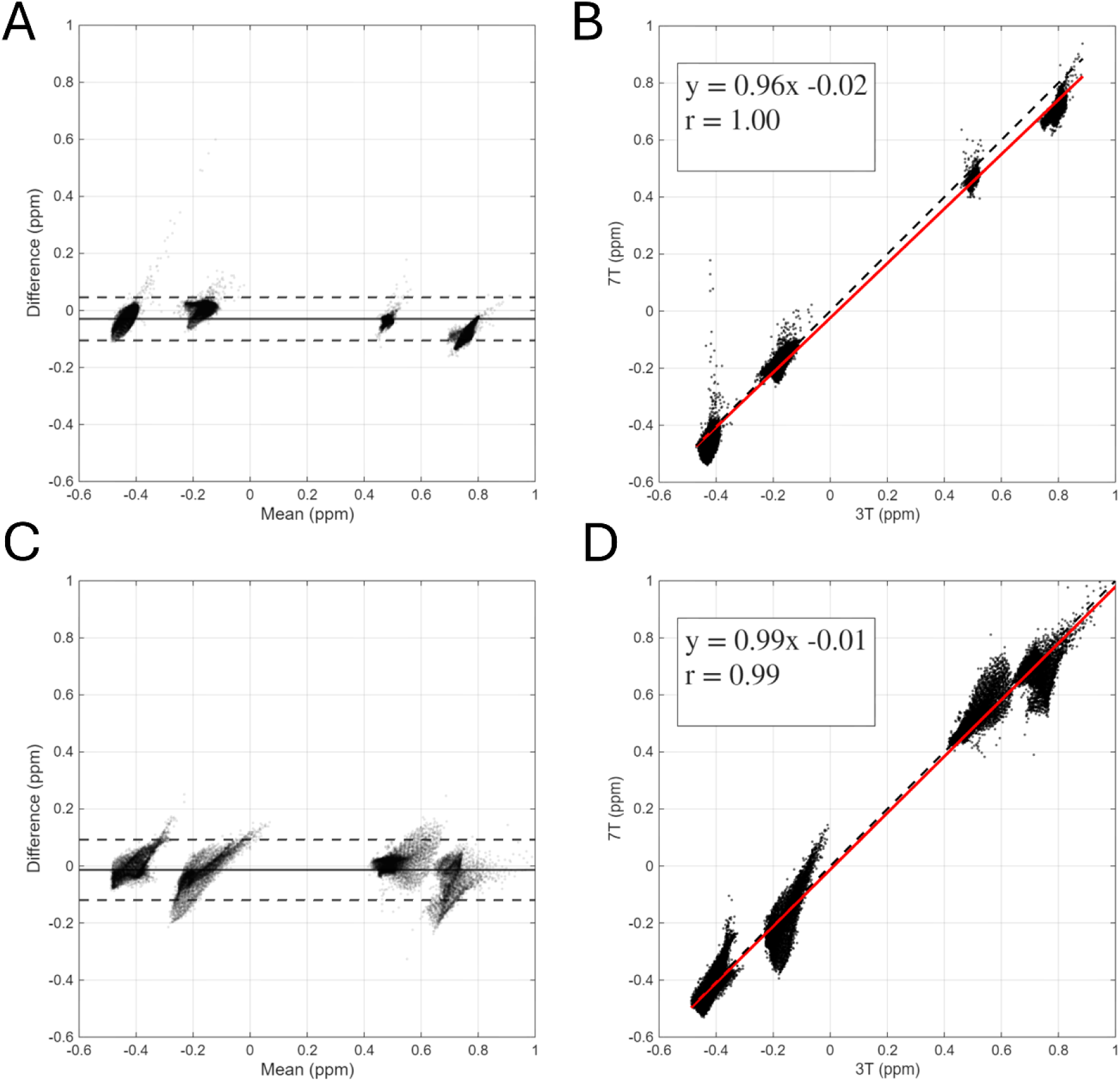
Quantitative comparison between 3T and 7T for MEDI (A-B) and COSMOS (C-D). For the Bland-Altman plots (A,C), the solid and dashed lines are the mean difference ± 2 × the standard deviation of the difference, respectively. For the correlation plots (B,D), the solid red lines indicate the trend lines in linear regression and the dashed line is the line of identity.

## 4. Discussion

### 4.1. QSM reconstruction

Our anthropomorphic phantoms were constructed from MR-compatible structural components, including nylon, medical amber resin and ABS filament. These structural components create signal voids, which are not typically observed in neurological data, since the cortical thickness is much less than the voxel size. These limitations were addressed by modifying two parameters in the QSM reconstruction. In our MEDI reconstructions, we modulated the weighting maps to reduce residuals deviating from a simulated mono-exponential signal – which we found reduced streaking artifacts at strong susceptibility interfaces. The weighting maps in Equation (4) were modulated using a threshold based on the statistics of the relative residual map (r_th_ = median(r) + 3·IQR(r)), which was roughly 10^2^ lower than the threshold used in neurological data (r_th_=0.3)^57^. We found that using the relative residual map statistics rather than a fixed threshold value gave better repeatability between orientations. Second, the gradient weighting mask in Equation (7) was derived by using a threshold (c∇) of 0.3, which is 1/3 of the threshold used in neurological data (c∇=0.9)^17^. Using too low c∇ resulted in streaking artifacts and image blurring, whereas using too high c∇ resulted in aliasing artifacts.

### 4.2. COSMOS-MEDI agreement

For the 4 ellipsoid ROIs (over a range of -0.6 ppm to 1.0 ppm): we observed excellent agreement between COSMOS and MEDI susceptibility at 3T, with linear regression of y=1.00x-0.01 (r=0.99). These observations were similar to the COSMOS and MEDI comparisons made by Liu et al. ^19^ who reported on a 3T scanner (over a range of -0.3 ppm to 0.5 ppm): y=0.97x-0.00 (r=0.97). At 7T, we observed a 7% underestimation of the MEDI relative to COSMOS. This is likely attributable to increased regularization bias at 7T and is reflected in ROI-based measurements for accuracy; the bias at 3T (-0.002 ppm) was lower than at 7T (-0.056 ppm), and the RMSE at 3T (0.082 ppm) was lower than at 7T (0.092 ppm). Higher field strength produces larger susceptibility-induced field gradients and greater intravoxel dephasing, which reduces the reliability of the local field map and increases the influence of MEDI’s morphology-based L1 regularization. The resulting edge-preserving regularization suppresses susceptibility amplitudes, particularly near sharp susceptibility interfaces, whereas COSMOS mitigates dipole inversion ill-conditioning through multi-orientation acquisition and therefore exhibits substantially less amplitude bias.

### 4.3. Crossfield-strength agreement

The motivation behind using an ultra-high-field scanner is to acquire images at increased spatial resolution without trading off SNR and imaging time. Optimizing and standardizing QSM protocols used for both clinical and ultra-high-field scanners are essential to make susceptibility a robust biomarker^65^. For the 4 ellipsoid ROIs (over a range of -0.6 ppm to 1.0 ppm): we observed excellent agreement between 3T and 7T susceptibility, with linear regression of y=0.96x-0.02 (r=1.00) for MEDI and y=0.99x-0.01 (r=0.99) for COSMOS. The mean difference (3T-7T) of -0.01 to -0.03 ppm mirrors observations in previous work^44^, where we observed mean differences of -0.01 ppm to -0.02 ppm using the same materials in a cylindrical phantom. Ferritin and CaCl_2_ were used as doping materials since each are linearly magnetizable, meaning that the magnetic susceptibility is invariant with magnetic field strength^44^.

### 4.4. Accuracy, precision and repeatability

The ellipsoids outperformed the straws in each of the 3 ROI-based metrics (accuracy, precision and repeatability). There are two reasons for this. First, the straw adapter was constructed from ABS, which led to strong susceptibility effects (see Figure 6A), which has been verified in another study (Δχ=2.748 ppm)^64^. Second, the volume covered along the straw’s radial axis (6 mm diameter) is much smaller compared to the ellipsoid’s minor axis (20.8 mm diameter). The sharp transitions in magnetic susceptibility at the edge boundaries led to phase aliasing and partial volume effects that make quantification difficult. From our experience using phantom, larger diameters (10 mm) make it easier to identify reliable voxels for susceptibility measurement.

The repeatability of the 7T MEDI susceptibility measurements (RC=0.459 ppm, RC’=0.141 ppm) outperformed the 3T MEDI susceptibility measurements (RC=0.652 ppm, RC’=0.168 ppm). The improved repeatability at 7T is likely due to the proximity of the receive coil elements to the phantom, leading to improved SNR across the imaging volume. The 3T acquisitions had much larger orientations about the x- and y-axes (see Tables S1-S2), leading to increased orientation-dependent variation. Signal void regions were associated with blooming artifacts occurring at phantom compartments containing nylon, ABS, air bubbles, and uncured resin sediments. Strong blooming artifacts in the complex GRE data appear near susceptibility interfaces orthogonal to the applied field vector^66^. These blooming artifacts can be reduced with smaller echo time; which is the subject of future work. The orientation-dependent local fields in MRI are a convolution of the dipole kernel with all susceptibility sources including non-local susceptibility sources^20,21^; which explains to some extent the imperfect repeatability of our susceptibility measurements.

### 4.5. Phantom design

This novel anthropomorphic phantom provided spheroids with thin walls (0.7 mm edge thickness) that theoretically reduces signal voids associated with phantom construction materials. Robust & thin-walled structures (spheroids) were 3D-printed and held together by a tapped thread (nylon bolts). From the susceptibility maps, the susceptibility of the (medical amber resin) ellipsoid walls appeared to be close to the susceptibility of the PVP40/NaCl background filling, limiting unintended susceptibility effects. However, other construction materials lead to stronger susceptibility effects (given relative to water): ABS (Δχ = 2.748 ppm)^64^, nylon (Δχ = -0.446 ppm)^42^, Araldite 2020 epoxy glue (Δχ = -0.662 ppm)^42^. Araldite 2020 epoxy glue was applied at the nylon screw threads due to loss of material (approximately 5 mm) during thread tapping. Moreover, since the doped agarose gel contracted after cooling, we were unable to prevent air (Δχ = 9.4 ppm) accumulating at the base of the fill port. From a practical perspective, assembling the nylon threads onto the jagged skull surface proved difficult, therefore future phantom studies might consider modification of the CAD file. By Curie’s Law, paramagnetic susceptibility is inversely proportional to temperature^63^; therefore, temperature should be regulated. Recent studies have used integrated temperature controls within the phantom, such as a thermostatic bath/circulator^67^ or an MR-visible liquid crystal thermometer^68^. This should be considered in future phantom design.

## 5. Conclusion

Herein, we developed an anthropomorphic phantom with various geometric compartments containing realistic susceptibilities. Reliable MEDI-based susceptibility measurements were obtained from ellipsoids but not from straws. The accuracy at 3T (bias = -0.002 ppm, RMSE = 0.082 ppm) was better than the accuracy at 7T (bias = -0.056 ppm, RMSE = 0.092 ppm). Using voxels from the 4 ellipsoid ROIs, we observed excellent agreement between COSMOS and MEDI susceptibility maps at 3T, with linear regression of y=1.00x-0.01 (r=0.99). We observed some underestimation of MEDI susceptibility maps relative to COSMOS at 7T, with linear regression and y=0.93x-0.04 (r=0.99). The results imply that QSM reconstructions are reliable with 3T scanners but can be challenging with 7T scanners at high magnetic susceptibilities.

## Acknowledgements

We are grateful to Massachusetts’ General Hospital (MGH) for providing the open-source CAD models of their head phantoms. We thank Martijn Cloos for providing the CAD model of the 7 Tesla Nova Medical 32 Rx-element coil. We thank John Steptoe from Queensland Brain Institute for 3D-printing support and engineering guidance. This research was funded by the Australian Government through the Australian Research Council (project number IC170100035). The authors acknowledge the facilities and scientific and technical assistance of the NIF, a National Collaborative Research Infrastructure Strategy (NCRIS) capability, at the University of Queensland.

## Data Accessibility Statement

We facilitate the reproducibility of this study and the phantom by providing imaging data (doi.org/10.5281/zenodo.21880376).

