## Supplementary Information for "A geometric anthropomorphic phantom for quantitative susceptibility mapping: accuracy and repeatability"

Table S1: Indexing of 3 T acquisitions with respective rotations and selection of acquisitions for COSMOS with their condition number.

| Index | 1(N) | 2 | 3 | 4 | 5 | 6 |
| --- | --- | --- | --- | --- | --- | --- |
| θ (^°^) | 0 | -10 | 11 | -40 | -4 | -9 |
| ψ (^°^) | 0 | -37 | -51 | 0 | 1 | 1 |
| A(θ, ψ) (^°^) | 0 | 38 | 52 | 40 | 4 | 9 |

Table S2: Indexing of 7 T acquisitions with respective rotations and selection of acquisitions for COSMOS with their condition number.

| Index | 1(N) | 2 | 3 | 4 | 5 | 6 |
| --- | --- | --- | --- | --- | --- | --- |
| θ (^°^) | 0 | -7 | -11 | -37 | -11 | -5 |
| ψ (^°^) | 0 | 23 | -22 | -3 | -19 | 21 |
| A(θ, ψ) (^°^) | 0 | 25 | 25 | 37 | 22 | 21 |

Table S3: ROI measurements from COSMOS-based susceptibility maps. χ_COSMOS,CF_ denoted closed-form solutions (Equation 8) and χ_COSMOS, IMPROVED_ denoted regularized solutions (Equation 9). All measurements given in ppm and reported as mean ± standard deviation.

|  | Ferritin 7.0 mM | Ferritin 10.2 mM | CaCl_2_ 0.9 M | CaCl_2_ 1.8 M |
| --- | --- | --- | --- | --- |
| χ_COSMOS, CF, 3T_ | 0.505 ± 0.123 | 0.779 ± 0.227 | -0.183 ± 0.213 | -0.494 ± 0.149 |
| χ_COSMOS, IMPROVED, 3T_ | 0.519 ± 0.052 | 0.806 ± 0.060 | -0.179 ± 0.047 | -0.474 ± 0.043 |
| χ_COSMOS, CF,_ _7T_ | 0.545 ± 0.101 | 0.755 ± 0.169 | -0.220 ± 0.143 | -0.489 ± 0.131 |
| χ_COSMOS, IMPROVED, 7T_ | 0.549 ± 0.069 | 0.757 ± 0.102 | -0.212 ± 0.069 | -0.474 ± 0.063 |

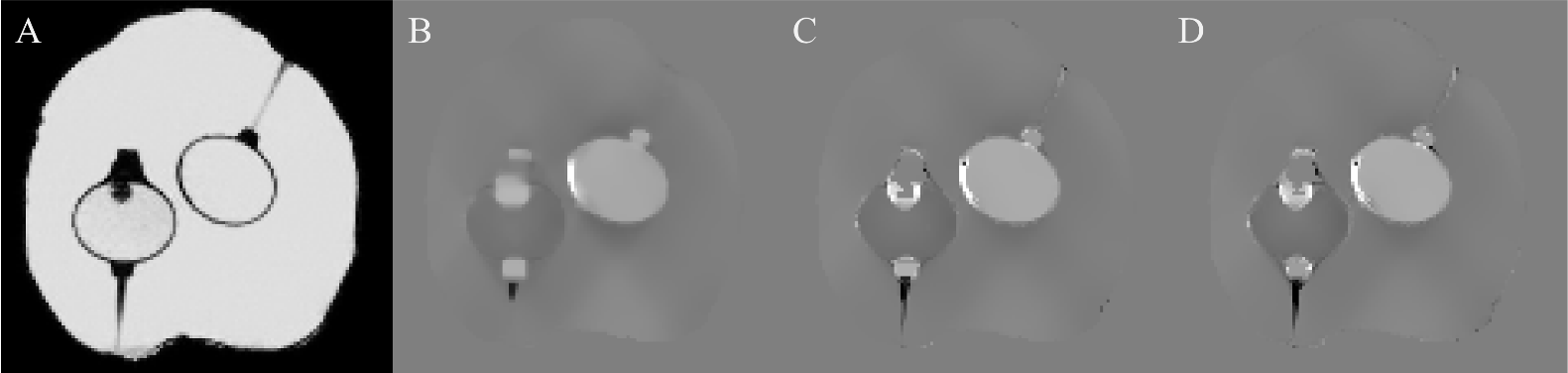

Figure S1: Magnitude image (A) and susceptibility maps (B-D) at varying levels of c_∇_: (B) c_∇_=0.1, (C) c_∇_=0.3, (D) c_∇_=0.5. Streaking artifacts (in the background filling) and image blurring were most prominent at c_∇_=0.1. Aliasing artifacts (at the edges of the ellipsoids) were most prominent at c_∇_=0.5. It was determined that c_∇_=0.3 was morphologically consistent with the magnitude image.

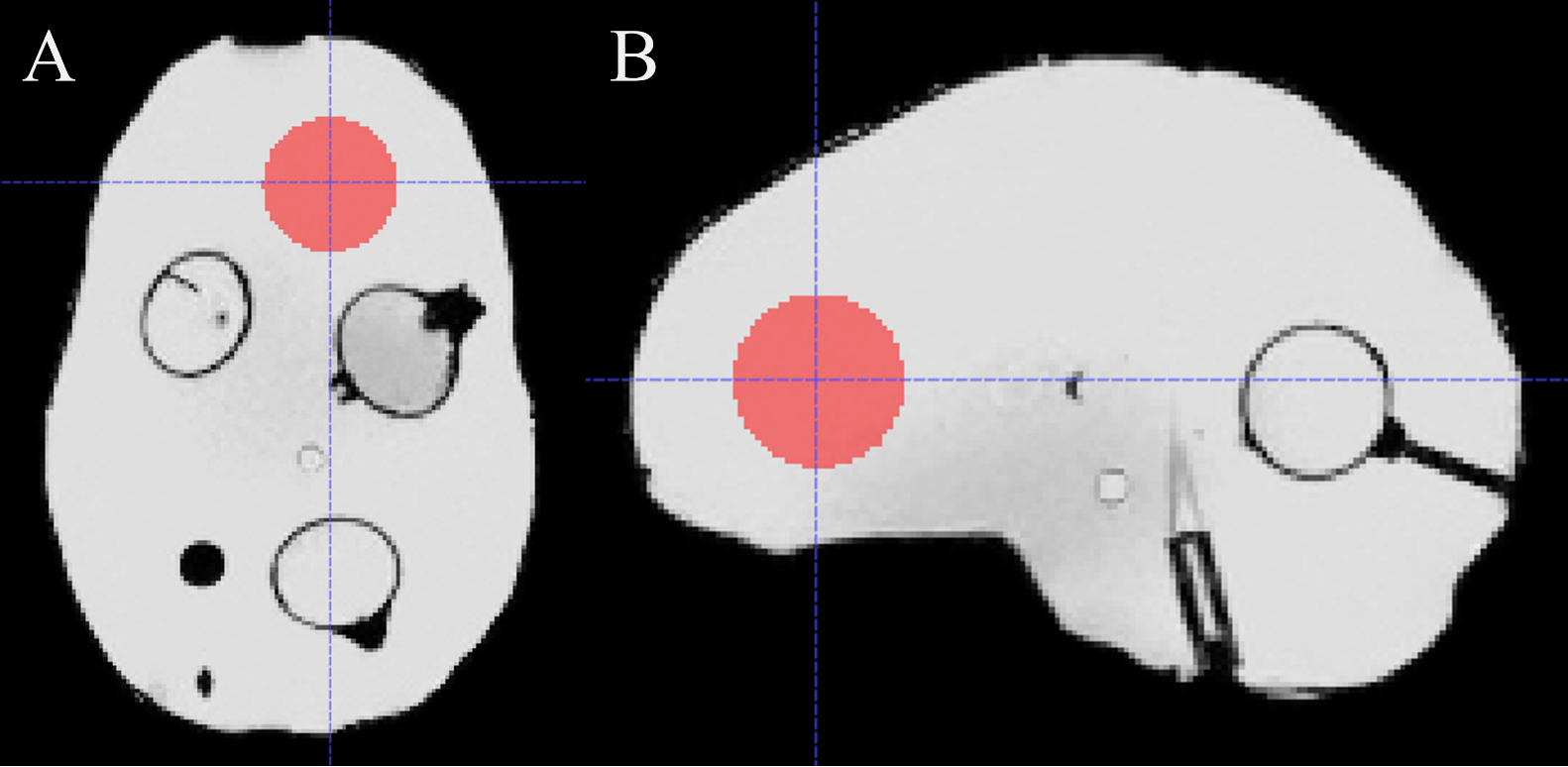

Figure S2: Reference mask (M_R_) shown in (A) coronal and (B) sagittal planes. M_R_ was obtained by defining a 15 mm sphere centered at the following coordinate (in mm): R∙[x_c_–10, y_c_+50, z_c_], where R denoted the 3×3 rotation matrix and [x_c_, y_c_, z_c_] the center-of-mass coordinates.

Figure S3: ROI-based susceptibility measurements plotted together with the literature values. Within-region standard deviation shown in vertical error bars.

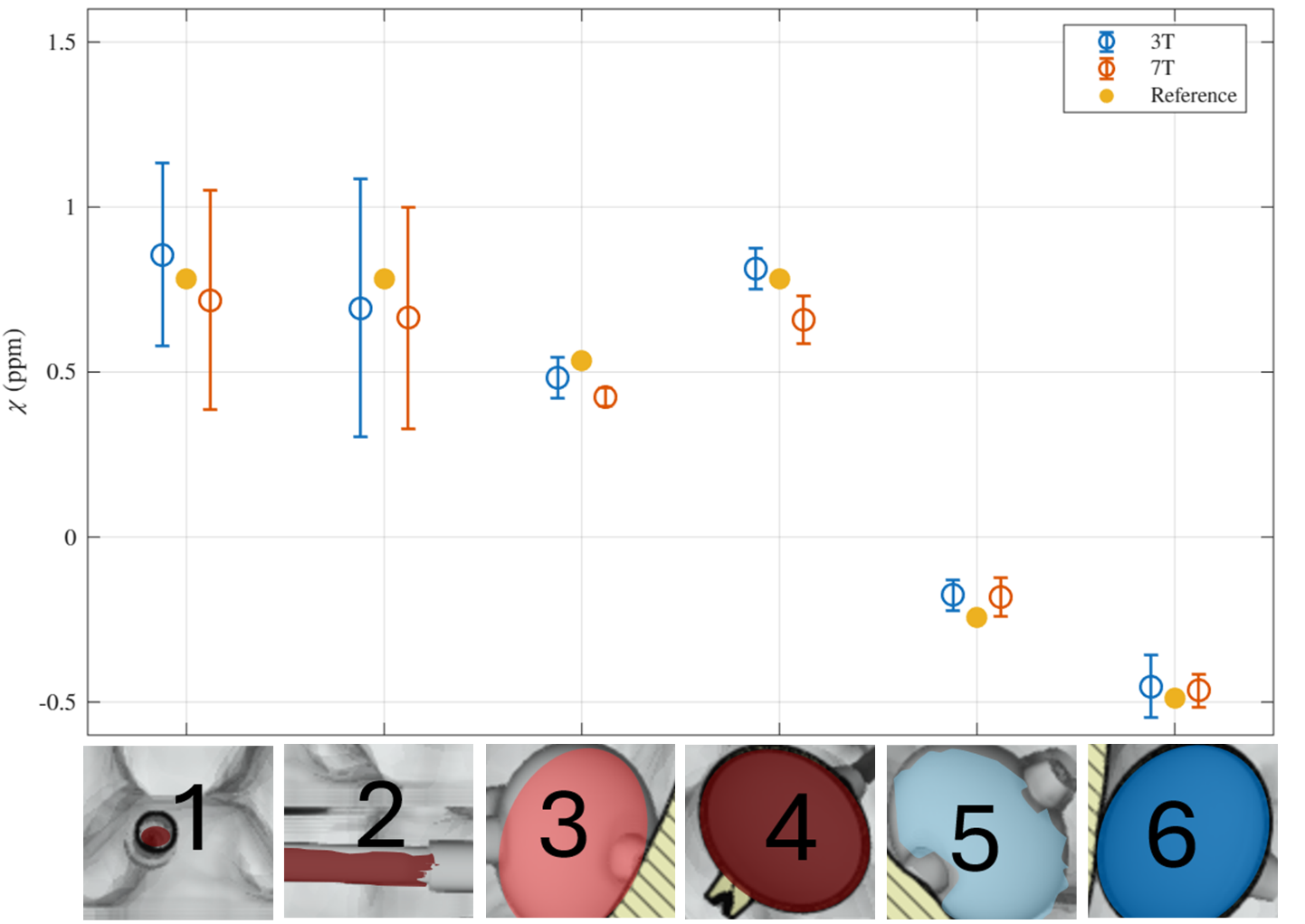

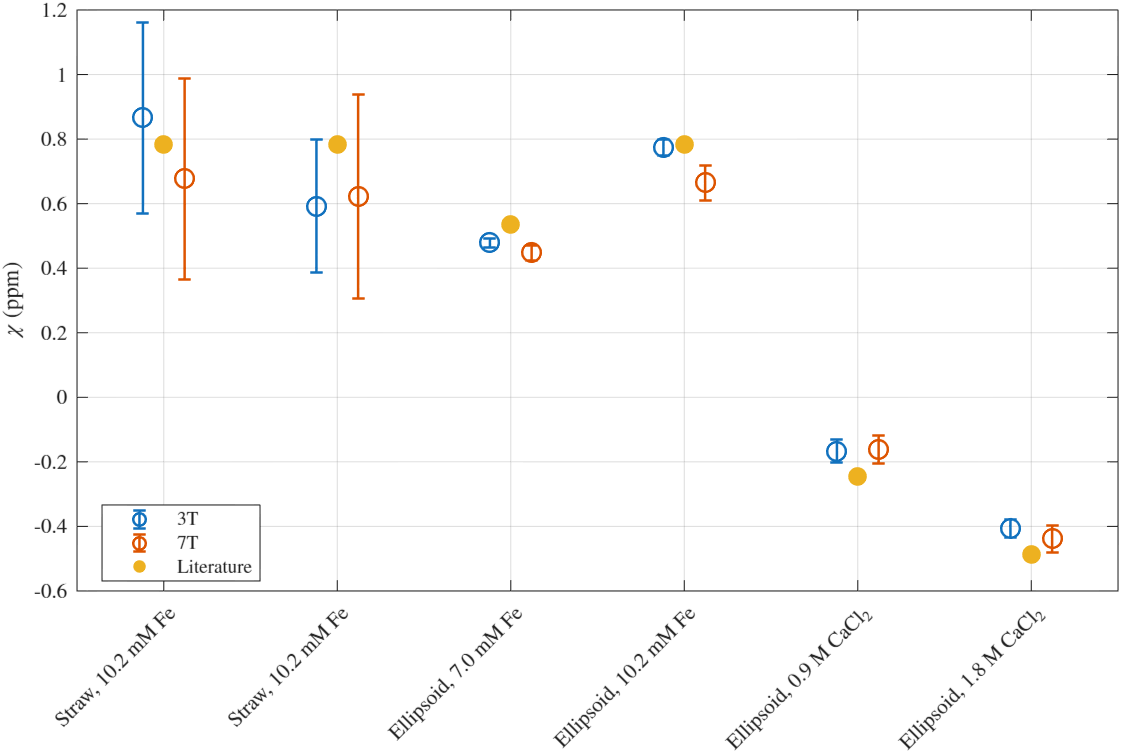
